# Reduced CREB3L2 expression is associated with resistance to sorafenib and poor prognosis in ER-positive breast cancer

**DOI:** 10.1101/078949

**Authors:** Sean Mullarkey, Vahid Arabkari, Muhammad Mosaraf Hossain, Sanjeev Gupta, Ananya Gupta

**Affiliations:** Discipline of Physiology, School of Medicine, National University of Ireland Galway, Ireland; Discipline of Pathology, School of Medicine, National University of Ireland Galway, Ireland

**Author notes:** Address correspondence to: Ananya Gupta, Discipline of Physiology, School of medicine, National University of Ireland Galway, Ireland.

**Keywords:** unfolded protein response, Sorafenib, CREB3L2, breast cancer

## Abstract

Sorafenib is a multikinase inhibitor that acts by inhibiting tumour growth and disrupting tumour microvasculature through anti-proliferative, anti-angiogenic, and pro-apoptotic effects. However, development of resistance to sorafenib often prevents its long-term efficacy. No validated biomarkers currently exist for appropriately selecting patients with cancer for sorafenib treatment. In the present study, we report that CREB3L2 expression in human breast cancer cell lines is a marker of response to sorafenib. Analysis of human breast cancer cell lines using Oncomine database and Genomics of Drug Sensitivity in Cancer (GDSC) database revealed an association between reduced expression of CREB3L2 and sensitivity to sorafenib. Wet lab experiment in five human breast cancer cell lines confirmed the association between reduced expression of CREB3L2 and sensitivity to sorafenib. Further, reduced expression of CREB3L2 was associated with poor Reccurence Free Survival in Luminal breast cancer. Our results suggest that CREB3L2 expression is a biomarker of response to sorafenib and outcome in breast cancer.

## INTRODUCTION

Sorafenib, (Nexavar) an oral multikinase inhibitor, possesses potential activity against several receptor tyrosine kinases including vascular endothelial growth factor receptor (VEGFR) 1, 2 and 3, as well as, platelet-derived growth factor receptor-β (PDGFR-β) [1, 2]. Sorafenib is approved by the U.S. Food and Drug Administration for the treatment of patients with advanced renal cell carcinoma (RCC) and hepatocellular carcinoma (HCC) [3, 4]. It is also approved by the European Medicines Agency for the treatment of patients with HCC and advanced RCC[4]. Preclinical studies suggest that sorafenib acts on tumours and tumour vasculature by inhibiting cellular proliferation and angiogenesis and/or by inducing apoptosis [5].

The CREB3L2 (also known as BBF2H7) is a novel endoplasmic reticulum (ER)-resident transcription factor that shares a region of high sequence similarity with ATF6. Like ATF6, CREB3L2 is activated through regulated intramembrane proteolysis (RIP) [6]. The CREB3L2 is the target of chromosomal translocation and produces chimeric oncoproteins in human cancer. In low grade fibromyxoid sarcoma (LGFMS), the bZIP and carboxyl domain of CREB3L2 is fused to N-terminal fragment of FUS [7]. FUS-CREB3L2 has a very high incidence (96%) in LGFMS and is a hyperactive transcription factor [7]. In thyroid follicular carcinoma, the N-terminal transactivation domain of CREB3L2 is fused to PPARγ [8]. The CREB3L2-PPARγ fusion protein can induce the growth of normal human thyroid cells by stimulating cell proliferation [8]. A recent RNAi screen revealed a role for CREB3L2 in the survival pathway in glioma cells [9]. The RAS-MAPK signalling cascade induces CREB3L2 expression, which directly upregulates ATF5 transcription. ATF5 then stimulates expression of the anti-apoptotic protein MCL1, which promotes cell survival [9]. Sorafenib treatment resulted in decreased cell viability of mouse malignant glioma and human malignant glioma cells that is associated with reduced expression of CREB3L2, ATF5 and MCL1 at both the protein and mRNA levels. However the role of CREB3L2 expression in determining the efficacy of sorafenib is not clear. Given the emerging role of CREB3L2 in human cancer we initiated further studies to interrogate its role in breast cancer.

## METHODS

### Cell culture and treatments

MCF7, T47D, MDA-MB231, BT474 and SKBR3 cells were purchased from ECACC. Cells were maintained in Dulbecco’s modified medium (DMEM) supplemented with 10% FCS, 100 U/ml penicillin and 100 mg/ml streptomycin at 37 °C with 5% CO_2_. All reagents were purchased from Sigma– Aldrich unless otherwise stated.

### Cell Viability Assay

Cells were seeded in 96-well at ~15% confluency in medium with 5% or 10% FBS and penicillin/streptomycin. One day following plating, cells were treated with range of concentrations of Sorafenib and then returned to the incubator for assay at a 72-hour time point. Cells were assayed with MTS reagent in accordance with the manufacturer’s instructions (Promega Corp., Madison, WI). In brief, 20 µL MTS reagent was added directly to the wells and cells incubated at 37°C for a maximum of 4 h. Assessment of metabolic activity was recorded as relative colorimetric changes measured at 490 nm. The data was transferred to Microsoft Excel and analysed. Background absorbance was corrected using triplicate sets of wells containing medium only (no cells) and MTS reagent as per experimental well. The results represent the mean ± SD of quadruple samples, expressed as a percentage of control. Each experiment was performed at least three times.

### RNA extraction, RT-PCR and real time RT-PCR

Total RNA was isolated using Trizol (Life Technologies) according to the manufacturer’s instructions. Reverse transcription (RT) was carried out with 2 μg RNA and random primers (Promega) using ImProm-II™ Reverse Transcription System (Promega). Real-time PCR method to determine the induction of UPR target genes has been described previously [10]. Briefly, cDNA products were mixed with 2 × TaqMan master mixes and 20 × TaqMan Gene Expression Assays (Applied Biosystems) and subjected to 40 cycles of PCR in StepOnePlus instrument (Applied Biosystems). Relative expression was evaluated using the ΔΔCT method.

### Statistical Analysis

The data is expressed as mean ± SD for three independent experiments. Differences between the treatment groups were assessed using Two-tailed paired student’s t-tests. The values with a p<0.05 were considered statistically significant.

## RESULTS and DISCUSSION

We examined the expression levels of CREB3L2 in a panel of 51 human breast cancer cell lines using the ONCOMINE Cancer Profiling Database (Fig 1). In the panel there were cell lines representing (27 cell lines) Basal-like, (10 cell lines) Luminal and (8 cell lines) HER2-amplified subtype. As shown in Fig 1, CREB3L2 expression levels showed a range of log2 median centered intensity from -2.8 to 1.6. The high proportion of Basal-like cell lines (62.9 %) showed CREB3L2 expression higher than the median while high proportion of Luminal cell lines (62.5%) showed CREB3L2 expression lower than the median. In contrast, HER2 amplified cell lines were equal distributed on either side of median. Next we analysed the sensitivity of the human breast cancer cell lines to sorafenib using the Genomics of Drug Sensitivity in Cancer (GDSC) database (www.cancerRxgene.org). As shown in Fig 2, the IC_50 value for sorafenib ranged from 0.279-537.37 μM in a panel of 29 breast cancer cell lines. We stratified the 29 breast cancer cell line into three categories: *Sensitive* (IC_50 < 20 μD), *Intermediate sensitivity* (IC_50 between 20-300 μD) and *Resistant* (IC_50 > 300 μD). We noticed that majority of cell lines from the *Resistant* group had CREB3L2 expression lower than the median while majority of cell line in *Sensitive* group had CREB3L2 expression higher than the median (Fig 3). These results suggested a putative association between CREB3L2 expression and response to sorafenib of human breast cancer cell lines. Next we performed experimental validation of predicted association between CREB3L2 expression and response to sorafenib. For this purpose we determined the expression of CREB3L2 in 5 breast cancer cell lines by qRT-PCR. As shown in Fig 4A, in agreement the ONCOMINE Cancer Profiling Database the expression of CREB3L2 was highest in MDA-MB231 and lowest in MCF7 cells. Further we evaluated the sensitivity of MDA-MB231 and MCF7 cells to sorafenib. We observed that in line with results of Genomics of Drug Sensitivity in Cancer (GDSC) database MCF7 cells were resistant to sorafenib treatment as compared to MDA-MB231 cells (Fig 4B). Taken together our results indicate an association between CREB3L2 expression and response to sorafenib of human breast cancer cell lines, where increased expression of CREB3L2 is a marker of response to sorafenib.

**Figure 1.**
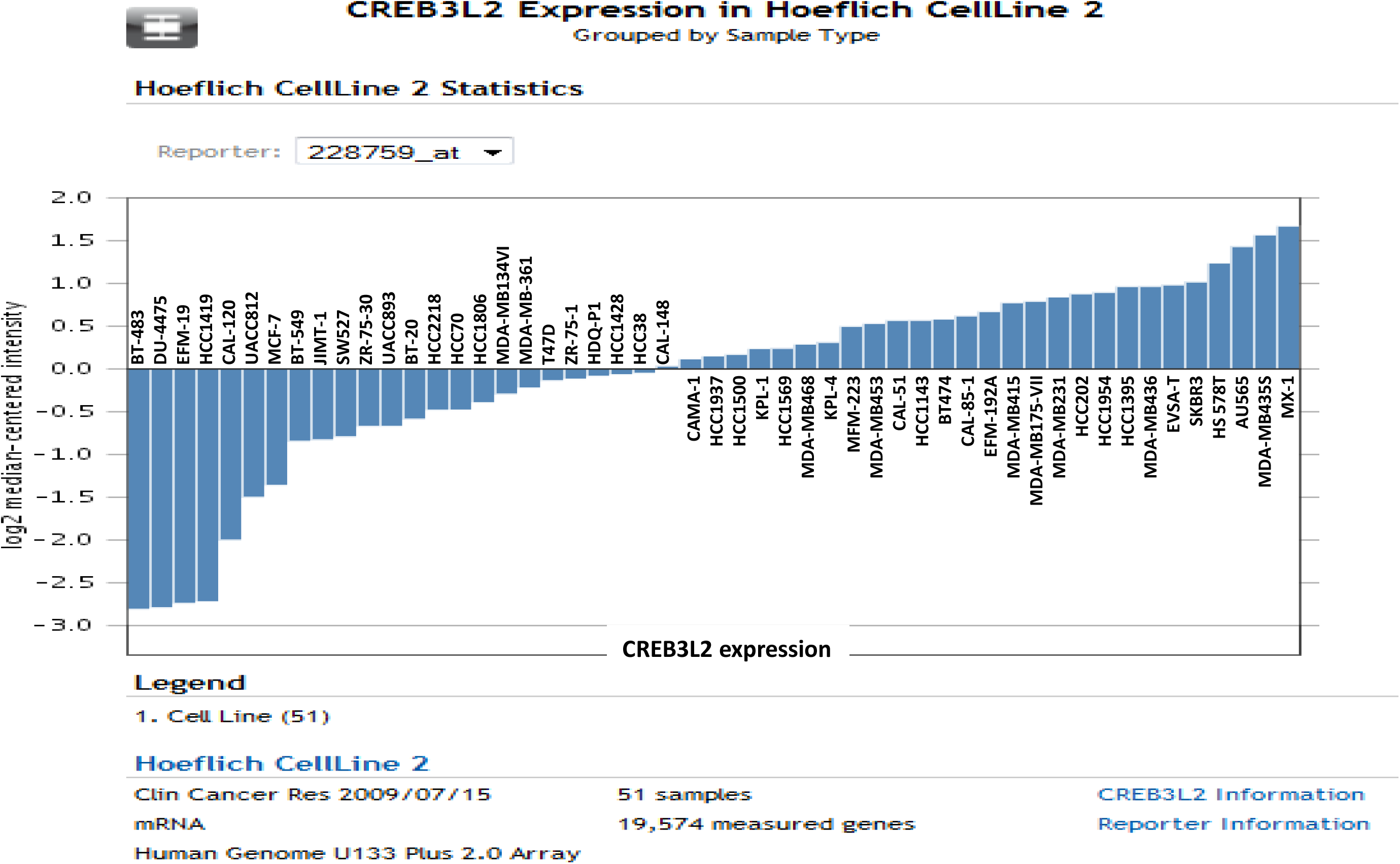
Expression of CREB3L2 in panel of human breast cancer cell lines. The Hoeflich cell line data was analysed using oncomine database for the expression of CREB3L2 in a panel of human breast cancer lines. The cell lines are arranged in the ascending order of CREB3L2 expression.

**Figure 2.**
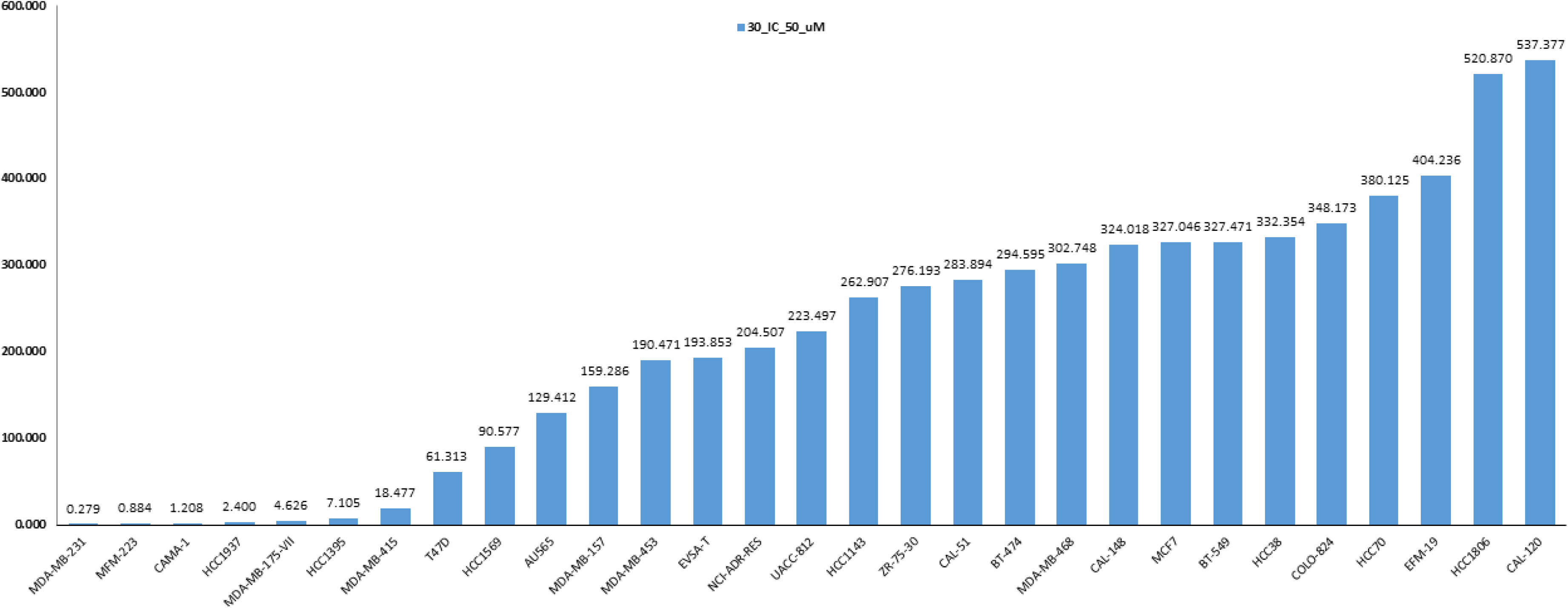
Sensitivity of human breast cancer cell lines to Sorafenib. Genomics of Drug Sensitivity in Cancer (GDSC) database (www.cancerRxgene.org) was analysed for the sensitivity to Sorafenib focusing on the human breast cancer cell lines. Cell lines are arranged in the ascending order of IC_50 value for sorafenib (μM).

**Figure 3.**
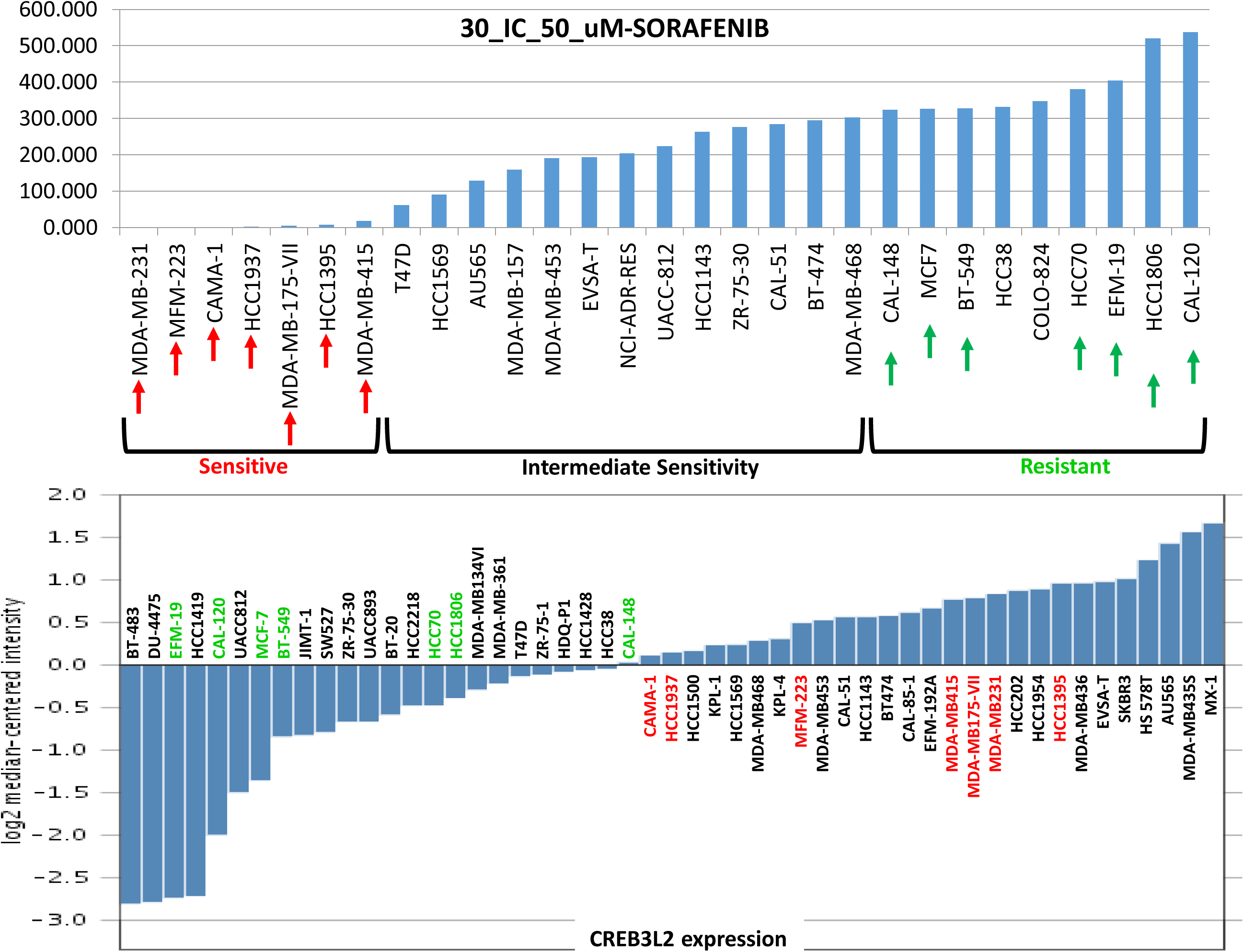
Putative association between CREB3L2 expression and sensitivity to sorafenib. *Upper panel,* human breast cancer cell lines are arranged in the ascending order of IC_50 value for sorafenib (μM). *Lower panel,* human breast cancer cell lines are arranged on the basis of CREB3L2 expression.

**Figure 4.**
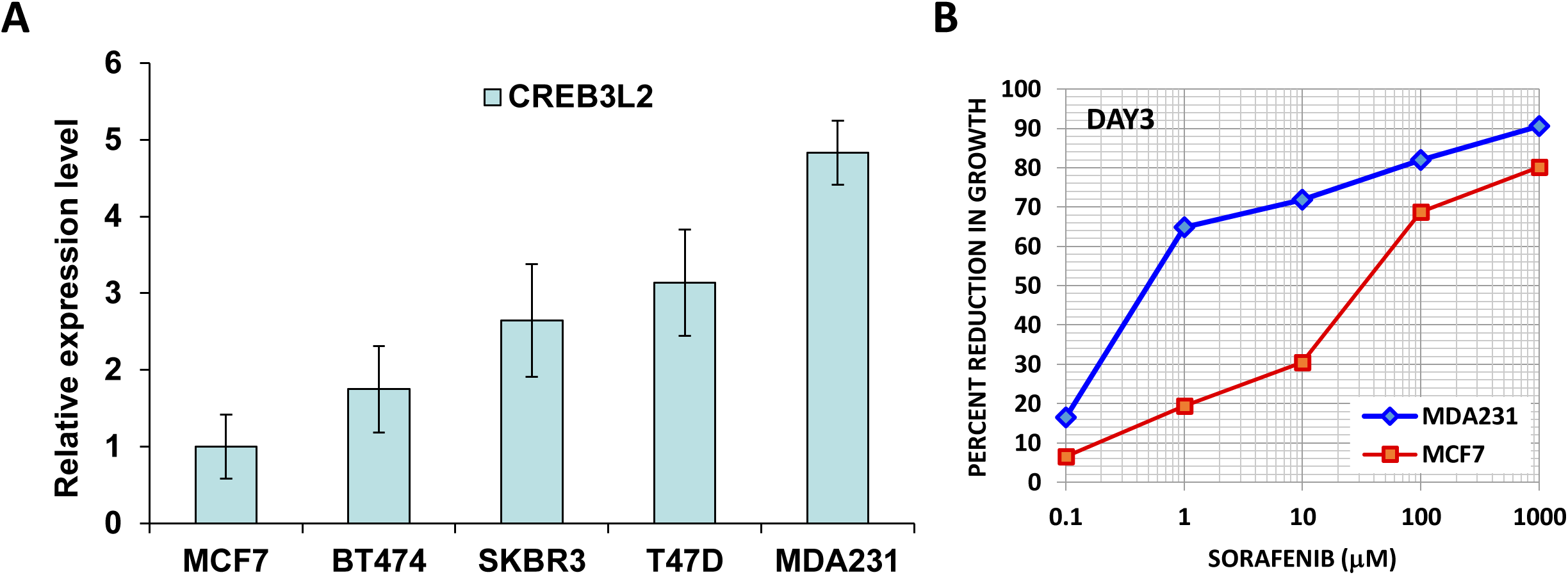
Association between CREB3L2 expression and sensitivity to sorafenib in human breast cancer cell lines. **(A)** Total RNA was isolated from the five indicated human breast cancer cell lines. The expression level of CREB3L2 was quantified by real-time RT-PCR, normalizing against RPLP0. **(B)** MCF7 and MDA-MB231 cells were treated with indicated doses of sorafenib for 72 hours. Line graphs show the percentage reduction in growth of the treated cells as compared to untreated control cells.

We evaluated the prognostic potential of CREB3L2 using web-based analysis tools (KM plotter and Breastmark) that uses microarray expression profiles and detailed clinical data to identify prognostic genes in breast cancer [11, 12]. Both algorithms predicted significant association of reduced CREB3L2 expression with poor recurrence free survival in ER-positive IBC (Fig 5). However the expression of CREB3L2 was not prognostic in HER2-amplified or Basal-like patients (Fig 5).

**Figure 5.**
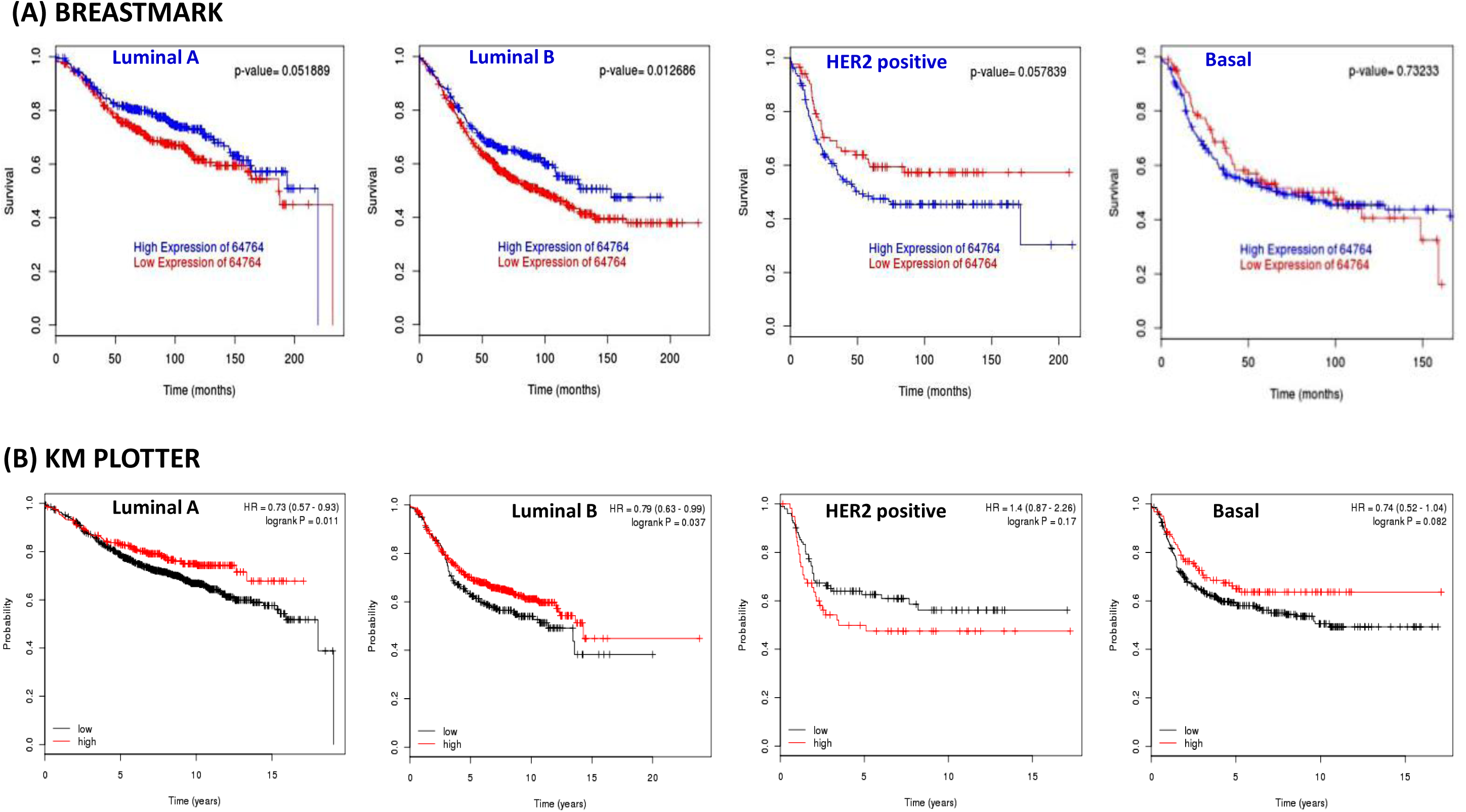
Kaplan-Meier curves in subtypes of invasive breast cancer for Relapse Free Survival (RFS) categorized according to CREB3L2 expression. Two web-based algorithms, (A) BreastMark (http://glados.ucd.ie/BreastMark/index.html) and (B) KM plotter (http://kmplot.com/analysis/) were used to evaluate the association between CREB3L2 expression with RFS. Both algorithms use the microarray gene expression data and detailed clinical data to correlate clinical outcome with differential gene expression levels.

In the present study, we demonstrated that CREB3L2 expression is dysregulated in breast cancer cell lines. We found that 62.5% of Basal-like cell lines had increased expression of CREB3L2. Thus, it is conceivable that CREB3L2 is involved in the growth of Basal-like cancers. However, the molecular mechanisms of dysregulated expression of CREB3L2 in breast cancer cells is still unclear. Tumour microenvironment stress has been shown to cause perturbation of ER functions. Specifically, hypoxia, nutrient limitation and low pH are known to activate the UPR in cancer microenvironment. These pathological changes in the tumour environment can adversely affect the folding and maturation of newly synthesized proteins, leading to ER stress, and induce expression of CREB3L2 in response to this ER stress in cancer cells. However, further investigation is needed to clarify what modulates the expression of CREB3L2 in cancer cells.

One of the crucial molecular pathways responsible for endocrine resistance in ER-positive breast cancer is the RAS/RAF/MAPK signalling cascade, where MAPK can physically interact with and activate ERα, contribute to endocrine resistance and has been associated with risk of relapse after adjuvant endocrine therapy [13]. It has been reported that, when added to endocrine therapy, sorafenib may complement the effect of anti-oestrogens in abrogating the ERα signalling and restoring tumour growth inhibition [14, 15]. Our results suggests that expression of CREB3L2 can be used to stratify the patients receiving sorafenib in such studies.

## ACKNOWLWDGEMENTS

We are grateful to the technical officers in Lambe Institute for Translational Research, NUI-Galway. S.M was a recipient of the CMNHS, NUI Galway summer research fellowship.

## AUTHOR CONTRIBUTION

The author(s) have made the following declarations about their contributions: Conceived and designed the experiments: AG and SG. Performed the experiments: AG, SM, and SG. Analysed the data and wrote the main manuscript text and prepared figures AG, SM and SG. All authors reviewed the manuscript.

## COMPETING INTERESTS

Authors declare no competing financial interests.

## REFERENCES

1. Moreno-Aspitia, A., Clinical overview of sorafenib in breast cancer. Future Oncol, 2010. 6(5): p. 655–63.

2. Gradishar, W.J., Sorafenib in locally advanced or metastatic breast cancer. Expert Opin Investig Drugs, 2012. 21(8): p. 1177–91.

3. Grandinetti, C.A. and B.R. Goldspiel, Sorafenib and sunitinib: novel targeted therapies for renal cell cancer. Pharmacotherapy, 2007. 27(8): p. 1125–44.

4. Lang, L., FDA approves sorafenib for patients with inoperable liver cancer. Gastroenterology, 2008. 134(2): p. 379.

5. Heravi, M., et al., Sorafenib in combination with ionizing radiation has a greater anti-tumour activity in a breast cancer model. Anticancer Drugs, 2012. 23(5): p. 525–33.

6. Kondo, S., et al., BBF2H7, a novel transmembrane bZIP transcription factor, is a new type of endoplasmic reticulum stress transducer. Mol Cell Biol, 2007. 27(5): p. 1716–29.

7. Panagopoulos, I., et al., Characterization of the native CREB3L2 transcription factor and the FUS/CREB3L2 chimera. Genes Chromosomes Cancer, 2007. 46(2): p. 181–91.

8. Lui, W.O., et al., CREB3L2-PPARgamma fusion mutation identifies a thyroid signaling pathway regulated by intramembrane proteolysis. Cancer Res, 2008. 68(17): p. 7156–64.

9. Sheng, Z., et al., A genome-wide RNA interference screen reveals an essential CREB3L2-ATF5-MCL1 survival pathway in malignant glioma with therapeutic implications. Nat Med, 2010. 16(6): p.671–7.

10. Gupta, A., D.E. Read, and S. Gupta, Assays for induction of the unfolded protein response and selective activation of the three major pathways. Methods Mol Biol, 2015. 1292: p. 19–38.

11. Gyorffy, B., et al., An online survival analysis tool to rapidly assess the effect of 22,277 genes on breast cancer prognosis using microarray data of 1,809 patients. Breast Cancer Res Treat, 2010. 123(3): p. 725–31.

12. Madden, S.F., et al., BreastMark: An Integrated Approach to Mining Publicly Available Transcriptomic Datasets Relating to Breast Cancer Outcome. Breast Cancer Res, 2013. 15(4): p. R52.

13. Bazzola, L., et al., Combination of letrozole, metronomic cyclophosphamide and sorafenib is well-tolerated and shows activity in patients with primary breast cancer. Br J Cancer, 2015. 112(1): p. 52–60.

14. Isaacs, C., et al., Phase I/II study of sorafenib with anastrozole in patients with hormone receptor positive aromatase inhibitor resistant metastatic breast cancer. Breast Cancer Res Treat, 2011. 125(1): p. 137–43.

15. Massarweh, S., et al., Impact of adding the multikinase inhibitor sorafenib to endocrine therapy in metastatic estrogen receptor-positive breast cancer. Future Oncol, 2014. 10(15): p. 2435–48.

